# The anterior cingulate cortex is involved in intero-exteroceptive integration for spatial image transformation of the self-body

**DOI:** 10.1101/2023.09.01.555872

**Authors:** Takafumi Sasaoka, Kenji Hirose, Toru Maekawa, Toshio Inui, Shigeto Yamawaki

## Abstract

Spatial image transformation of the self-body is a fundamental function of visual perspective-taking. Recent research underscores the significance of integration of intero-exteroceptive information to construct representations of our embodied self. This raises the intriguing hypothesis that interoceptive processing might be involved in the spatial image transformation of our self-body. To test this hypothesis, the present study used functional magnetic resonance imaging to measure brain activity during an arm laterality judgment (ALJ) task. In this task, participants were tasked with discerning whether the outstretched arm of a human figure, viewed from the front or back, was the right or left hand. The reaction times for the ALJ task proved longer when the stimulus presented orientations of 0°, 90°, and 270° relative to the upright orientation, and when the front view presented as compared to the back view. Corresponding to the increased reaction time, increased brain activity was manifested in a cluster centered on the dorsal anterior cingulate cortex (ACC). Furthermore, this cluster of brain activity exhibited overlap with regions where the difference in activation between the front and back views positively correlated with the participants’ interoceptive sensitivity, as assessed through the heartbeat detection task, within the pregenual ACC. These results suggest that the ACC plays an important role in integrating intero-exteroceptive cues for the purpose of spatially transforming the image of our self-body.

## 1. Introduction

Visual imagery can also be generated from different perspectives, highlighting a crucial cognitive function known as visual perspective-taking (VPT). This ability plays a pivotal role in estimating the mental states of others during social communication, as stated by researchers Frith and Frith (2006). To achieve this, brain engages in spatial image transformation of the body, as exemplified in the work of Kessler and Thomson (2010). Research focusing on the sense of self has indicated that neural representations of the body, including interoception ―our sense of internal body states―contribute significantly to shaping our sense of self (Seth, 2013). For example, studies have found intriguing connections between bodily perceptions and self-representation. The extent of proprioceptive drift in the rubber hand illusion (RHI) has been inversely linked to the interoceptive accuracy measured through a heartbeat counting task (Tsakiris et al., 2011). This suggests a meaningful interplay between interoception and self-representation. Moreover, there is evidence of a connection between autonomic responses and motor imagery (Ohishi et al., 2000; Ohishi and Maeshima, 2004), implying that our physical simulations trigger internal changes too. Therefore, the process of transforming our self-body’s spatial imagery likely involves the integration of both exteroception and interoception,

However, direct evidence directly linking interoceptive processing to the spatial image transformation of the body is currently absent. A previous study attempted to bridge this gap by examining the connection between performance on the VPT tasks and interoceptive accuracy, assessed using the heartbeat counting task (Erle, 2019). The results showed a negative correlation between indices of participants’ VPT difficulty, such as reaction times, and their interoceptive accuracy from the heartbeat counting task. This suggests that those with lower the interoceptive accuracy tended to perform less effectively in VPT tasks, reinforcing the relationship between VPT and interoception.

Moreover, the neural mechanisms underpinning integration of interoception and exteroception for the spatial image transformation of the self-body is unknown. Although some neuroimaging studies have delved into brain activity during the spatial transformation of one’s own body (Ganesh et al., 2015) and hand (de Lange et al., 2006), the involvement of interoceptive processing to these spatial image transformations remains a gap in our understanding. Should interoceptive processing indeed play a role in the spatial image transformation of the self-body, it would be reasonable to anticipate the activation of brain regions associated with intero-exteroceptive integration, such as the anterior insula and anterior cingulate cortex (ACC), during tasks necessitating this transformation. These areas, components of the salience network (Seeley et al., 2007) represent the highest order of interoceptive processing (Critchley and Harrison 2013; Savitz and Harrison 2018). They have also been posited as pivotal in the emergence of emotions and the embodied self (Seth, 2013). Thus, these critical regions governing interoceptive processing might similarly contribute to the spatial image transformation of the self-body.

Therefore, in the present study, we aimed to clarify the intero-exteroceptive integration of spatial image transformation of the self-body and its neural basis. To this end, we used a mental image transformation task in which participants performed arm laterality judgment (ALJ) concerning human figures presented in front or back views, rotated across various orientations (Parsons, 1987). Employing functional magnetic resonance imaging (fMRI), we monitored the participants’ brain activity during the ALJ task aiming to substantiate the following hypotheses:

H1: If interoceptive processing is involved in ALJ, then performance on the ALJ task should correlate with individual differences in interoception.

H2: If ALJ is reliant on intero-exteroceptive integration, brain regions associated with exteroceptive (e.g., visual) imagery transformation, such as the parietal lobe and premotor cortex (Zacks, 2008) along with regions linked to interoceptive processing (e.g., anterior insula and ACC) should show associations with ALJ task performance.

H3: Activity in brain regions responsible for integrating intero-exteroceptive information (e.g., anterior insula and ACC) during the ALJ task should correlate with individual differences in interoception.

To assess individual differences in interoception, we utilized a heartbeat discrimination task (HDT); (Whitehead et al., 1977; Maekawa et al., 2021), wherein participants assessed whether beep sounds, occurring at various delays from their R-peak, synchronized with their own heartbeat. In this study, we examined whether HDT-derived indices were related to behavioral ALJ performance and brain activation during the ALJ task.

## 2. Materials and Methods

### 2.1. Participants

Thirty-six undergraduate and graduate students took part in the experiment (9 women, age 21.7 ± 3 years, all were right-handed with a laterality coefficient of 0.93 ± 0.9). According to self-reported data, none of the participants had a history of mental disorders. Prior to participating in the study, all participants provided written informed consent in accordance with the principles of the Declaration of Helsinki. The Research Ethics Committee of Hiroshima University approved this study (approval number: E-965).

### 2.2. Arm laterality judgment task

#### 2.2.1. Stimuli

We used line drawings depicting either front or back views of a human body, featuring an outstretched right or left arm. These drawings were presented in four orientations:0°, 90°, 180°, and 270° from an upright position, encompassing a clockwise rotation on the image plane. In total, there were 16 distinct types of stimuli (Figure 1a). These visual stimuli were presented through a pair of MRI-compatible liquid crystal displays (Visual System HD; Nordic Neurolab, Norway).

**Figure 1.**
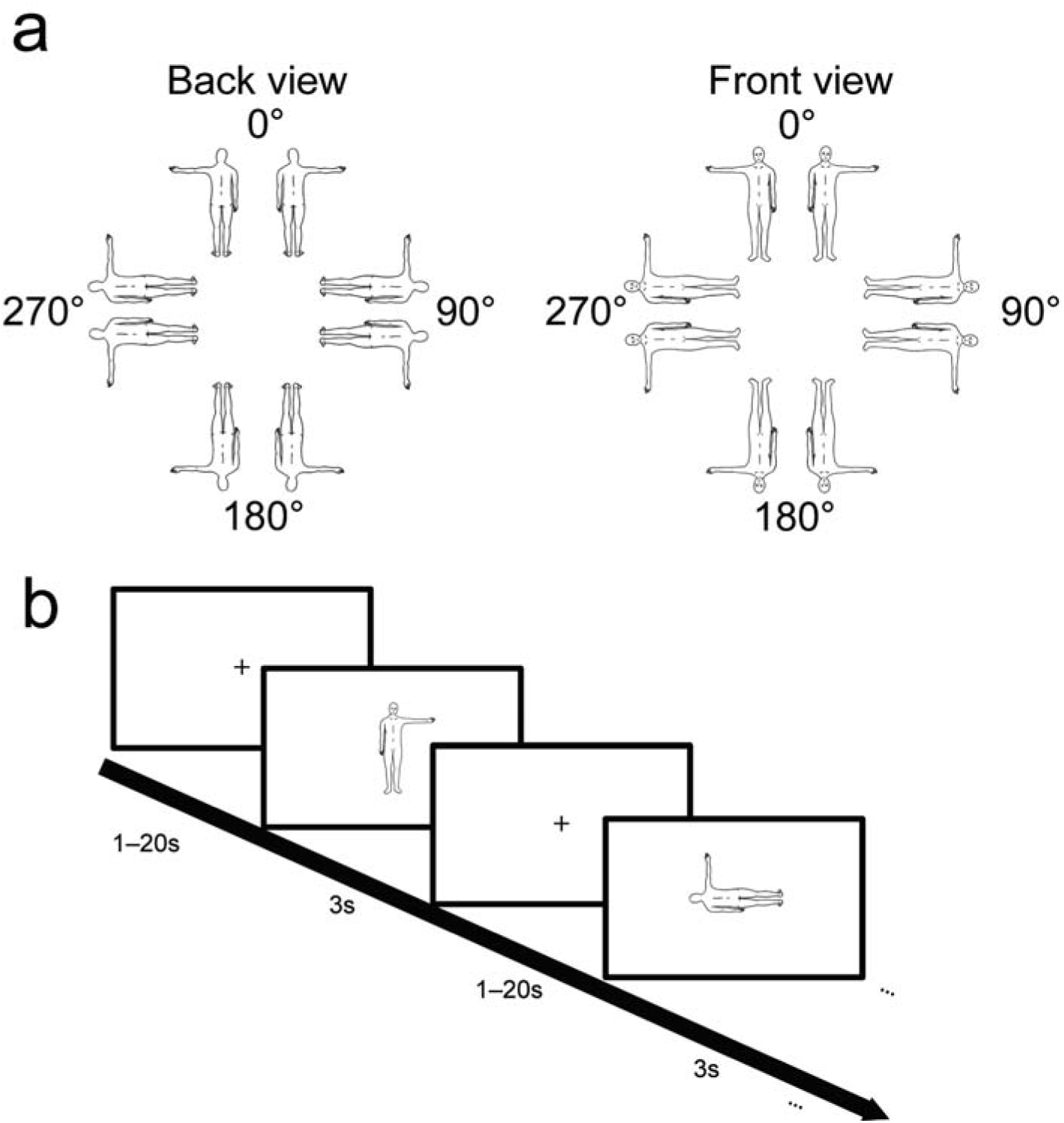
Procedure of the arm laterality judgment (ALJ) task. (A) The stimuli used in the task. (B) The timeline of trials of ALJ task. A stimulus was presented for a duration of 3 s, followed by an inter-stimulus interval (ISI) ranging from 1–20 s. During the ISI, a fixation cross was displayed. Refer to the main text for the details of the procedure.

#### 2.2.2. Task procedure

During each trial, a stimulus was displayed for 3 s. Participants were instructed to promptly and accurately indicate whether the outstretched arm belonged to the right or left hand. This response was obtained using an MRI-compatible foot pedal (Current Designs, Inc., Philadelphia, PA, USA), with each foot pedal corresponding to a specific hand. Each stimulus type was presented 20 times, resulting in 320 trials. (Current Designs, Inc., Philadelphia, PA, USA) with their feet as quickly and accurately as possible (20 trials per stimulus). The period between successive stimuli, referred to as the interstimulus interval (ISI), was set within the range of 1–20 seconds. During the ISI, participants were presented with a fixation cross and were asked to maintain their focus on it (Figure 1b). To optimize the timing of stimulus presentations in accordance with an event-related fMRI design, we used the OptSeq2 software (Dale, 1999). Should a participant fail to respond within three seconds, the trial concluded, and the next trial commenced. The 320 trials were distributed across four experimental runs, with each run consisting of 80 trials. Within each run, participants engaged in five trials for each of the 16 different stimulus types.

### 2.3. Heartbeat discrimination task

Outside the MRI scanner environment, participants performed a HDT in which they were presented with beep sounds with varying delays in relation to their R-peak detected by an electrocardiogram (ECG). We adopted the procedure with four delay conditions as outline by Maekawa et al. (2021). A three-lead ECG was attached to the chest of each participant. The ECG signal was recorded using a Biopac MP-160 system at a sampling rate of 1,000 Hz (Biopac Systems, Goleta, CA, United States). This signal was monitored using MATLAB 2020b (MathWorks, Inc., Natick, MA, United States) via a Biopac hardware application programming interface (BHAPI 2.2). When the R-peak was detected, a beep sound (1,000 Hz sine wave) was presented with various delays in the timing of the R-peak. Although there were some delays between the timings of R-peak detection and sound presentation owing to mechanical factors, the delay was within approximately 50 ms. There were four distinct delay conditions: 0, 150, 300, and 450 ms. Within each trial, a set of ten beep sounds were presented. Participants were tasked with determining whether the timing of the beep matched with the timing of the heartbeat and to subsequently convey their confidence level regarding the judgment. Six trials were conducted for each delay condition, totaling 24 trials.

### 2.4. Behavioral data analysis

#### 2.4.1. Arm laterality judgment task

In assessing the mean correct reaction time (RT), we conducted a two-way repeated-measures analysis of variance (rmANOVA) with front/back and orientation factors (0°, 90°, 180°, and 270°). Multiple comparisons were conducted using Bonferroni correction. For each front/back and orientation condition, trials in which stimuli were presented with both the right and left arms outstretched were collapsed. To exclude outliers, the mean and standard deviation (SD) of the correct RTs were computed for each run. We excluded the correct RTs falling beyond the mean ±3 SD from the analyses.

#### 2.4.2. Heartbeat discrimination task

Adopting the analytical approach devised by Maekawa et al. (2021), we employed a Gaussian function fitting procedure to the model participants’ responses. The resulting parameters of the function were subsequently regarded as indices reflecting individual differences in interoception. Considering the inherent periodicity of the heartbeat, we introduced an extra data point by considering the 450 ms condition as one obtained at (450 - mean R-R interval) (RRI) for each participant. Five data points (450 - RRI ms, 0 ms, 150 ms, 300 ms, and 450 ms) were fitted using a Gaussian function defined by the following equation:

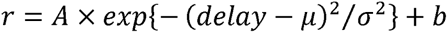

where *r* represents the ratio of the number of “matched” responses within a given delay condition to the number of trials (six in the present study). The variable *delay* corresponds to the delay conditions, *A*, μ, σ, and *b* stand for fitting parameters and represent the amplitude, center, width, and bias of the Gaussian function, respectively. Among these parameters, two are particularly pertinent as indicators of individual differences in interoception: (i) The amplitude of Gaussian (*A*) signifies the degree of consistency with which a participant produced “matched” responses in a specific delay condition compared to others, (ii) the width of the Gaussian (σ) signifies the range of delays encompassing participants’ “matched” responses. Maekawa et al. (2021) suggested that the *A* corresponds to interoceptive sensitivity (IS), and that the σ to inverse of interoceptive accuracy. Maekawa et al. (2021) demonstrated a positive association between IS and physiological responses (heart rate changes) triggered by musical stimuli and brain activation in the mid-insula. Based on their study, we focused on the amplitude of Gaussian (*A*) (hereafter, we refer *A* to IS following Maekawa et al. (2021)) and conducted the correlation analysis between the IS and the ratio of the correct RTs for front view stimuli relative to those for the back view stimuli. This choice stemmed from the expectation that the ALJ task for front views would involve spatial image transformations of the self-body, such as the mental simulation of “looking-back,” unlike that for back views. To assess the normality of the IS distribution (fitted parameter A), we conducted a Kolmogorov-Smirnov test to evaluate null hypothesis that IS follows a normal distribution. Our findings indicated a rejection of the null hypothesis, yielding a *p* value of 0.018. Therefore, we employed Spearman’s rank correlation coefficients for our correlation analysis. For this purpose, we used a robust correlation toolbox (Pernet et al., 2013) running on MATLAB 2022b.

### 2.5. MRI data acquisition

Anatomical and functional images were obtained using a 3.0 T MRI (Siemens MAGNETOM Skyra, Siemens AG, Munich, Germany). The T2*-weighted functional images were obtained with the following parameters: TR= 1,000 ms, TE = 30 ms, flip angle = 80°, field of view = 192 mm, 42 slices, voxel size of 3 × 3 × 4 mm (with 25 % gap), and a multiband factor of 3. For the T1 weighted anatomical images, a magnetization-prepared rapid gradient-echo imaging sequence with the following parameters: TR = 2,300 ms, TE = 2.98 ms, voxel size of 1 × 1 × 1 mm; flip angle of 9°; and field of view of 256 mm.

### 2.6. MRI data analysis

MRI data were analyzed using SPM12 software (Wellcome Department of Cognitive Neurology, London, United Kingdom; www.fil.ion.ucl.ac.uk/spm). A total of 734 echo-planar images (EPIs) were obtained for each run. To account for T1 equilibration effects, the initial ten EPIs were disregarded. The remaining 724 EPIs were used for preprocessing steps. The EPIs were realigned with the first EPI of each respective run. The T1 weighted anatomical images were then coregistered with the mean functional image, calculated using the EPIs from each run. The normalization procedure employed involved the alignment of the T1 images and realigned EPIs using the Montreal Neurological Institute (MNI) brain template. Following normalization, the EPIs underwent an 8-mm Gaussian kernel smoothing, resulting in a full width at half maximum of 8 mm.

For the first-level analysis, the preprocessed EPIs were employed. We defined the presentation of the 16 stimulus types, foot responses, and ISIs as events of interest. Each event was modeled using a boxcar function, factoring in the event’s onset and duration. We defined the durations of the stimulus presentation and foot responses as zero and the durations of the ISIs as their specific lengths. These event-related time series were then convolved with a canonical hemodynamic response function (HRF) and incorporated into a general linear model (GLM). Furthermore, the time series generated from the six head movement parameters, derived from the realignment process, were included in the GLM to account for head movement effects. To mitigate low-frequency noise, a high-pass filter with a cut-off period of 128-s was applied. Additionally, artifacts resulting from autocorrelation were corrected using a first-order autoregressive model. In the context of the first-level analysis, the regression coefficients for each regressor were estimated by the GLM uniquely tailored for each participant.

For the exploration of brain activity associated with the presentation of each stimulus type, contrast images were generated for each regressor. To assess the distinctions in brain activities between front and back views, corresponding contrast images were created. These contrast images subsequently served as inputs for group-level analysis. A one-sample *t*-test was conducted to identify significant brain activation for each event. To examine the main effects stemming from the front/back and orientation and their interaction on brain activation during stimulus presentation, a two-way rmANOVA was executed, including front/back and orientation factors. The reported brain regions exhibiting activation met the threshold of an uncorrected *p* < 0.001 at the voxel level, with a family wise error (FWE) corrected for multiple comparisons at *p* < 0.05 at the cluster level. To examine the effect of IS on the brain regions involved in ALJ, we conducted a multiple regression analysis using IS (the fitted parameter A) of each participant derived from the HDT as a covariate.

### 2.7. Generalized psycho-physiological interaction analysis

To examine ALJ-related functional connectivity (FC) and the effect of individual interoception differences on FC, we conducted a generalized psychophysiological interaction (gPPI) analysis through the use of the CONN toolbox (version 21.a; Whitfield-Gabrieli and Neto-Castanon, 2012). Based on the results that the activation in the cluster including the dorsal ACC (dACC) showed an interaction between the front/back and orientation factors and that this cluster and the brain region were positively associated with the IS overlapped in the pregenual ACC (pgACC; See Results section), we determined the bilateral pgACC to be the seed region. To define the regions of interest (ROI) in the pgACC, we referred to the anatomical ROIs based on the BrainNetome atlas (Fan et al., 2016).

For the gPPI analysis, the GLMs for each participant were imported into the CONN toolbox. For each participant, the gPPI was calculated using a multiple regression model, for the blood oxygenation level-dependent (BOLD) signal time series at each voxel. The regression model was characterized by three separate models: the main psychological factor, that is, all task effects convolved with a canonical HRF; the main physiological factor, that is, the BOLD time series in the seed regions; and the PPI term, which is the interaction term specified as these products. By calculating gPPI maps featuring the regression coefficients of the PPI terms for each ROI, we were able to derive the resulting gPPI maps. These gPPI maps were subjected into group-level analysis, facilitating the examination of both ALJ-related- and interoception-related FC through one-sample *t*-tests. As for the ALJ-related FC, we explored the brain regions showing FC with the pgACC when front views were presented. As for the interoception-related FC, we explored the brain regions showing FC with the pgACC correlated with the IS of each participant when the front view in the upright orientation was presented, based on the result that we observed a positive correlation between the IS and ALJ performance in the upright orientation.

## 3. Results

### 3.1. Behavioral data

#### 3.1.1. Correct reaction times for each stimulus

A two-way rmANOVA with front/back and orientation factors revealed a main effect of front/back (*F* (1,35) = 91.89, *p* < 0.001, partial η^2^ = 0.724), a main effect of orientation (*F* (1.707, 59.760) = 81.15, *p*_corrected_ < 0.001, partial η^2^ = 0.699, Greenhouse-Geisser correction was applied to control for violation of the sphericity assumption), and an interaction (*F* (2.089, 73.100) = 41.50, *p*_corrected_ < 0.001, partial η^2^ = 0.542) (Figure 2). Following this significant interaction, further analysis was conducted to ascertain the simple main effects for each orientation and front/back view. The simple main effects of the front/back factor were found to be significant for all orientations (*F*s (1, 35) >4.712, *p*s < 0.05, partial η^2^s > 0.119). Multiple comparisons revealed that for the 0°, 90°, and 270° orientations, RTs were longer for the front view compared to the back view (*p*s < 0.001). For the 180° view, the RTs for the back view were longer than those for the front view (*p* < 0.05). The simple main effects of orientation on both front and back views were also significant (*F*s (3, 33) > 4.772, *p*s < 0.01, partial η^2^ > 0.303). Multiple comparisons revealed that the RTs for the front view was longer for 180° than for 0°, 90°, and 270° (0°–90°, 0°–180°, 0°–270°, 90°–180°, 180°–270°: *p*s < 0.05; other pairs: *p*s > 0.708). For the back view, differences were significant for all pairs except for the pair between 90°and 270°, (90°–270°: *p* = 1.000; other pairs: *p*s < 0.001), indicating that the RTs for the back view were longer in the order of 180°, 90°/270°, and 0°.

**Figure 2.**
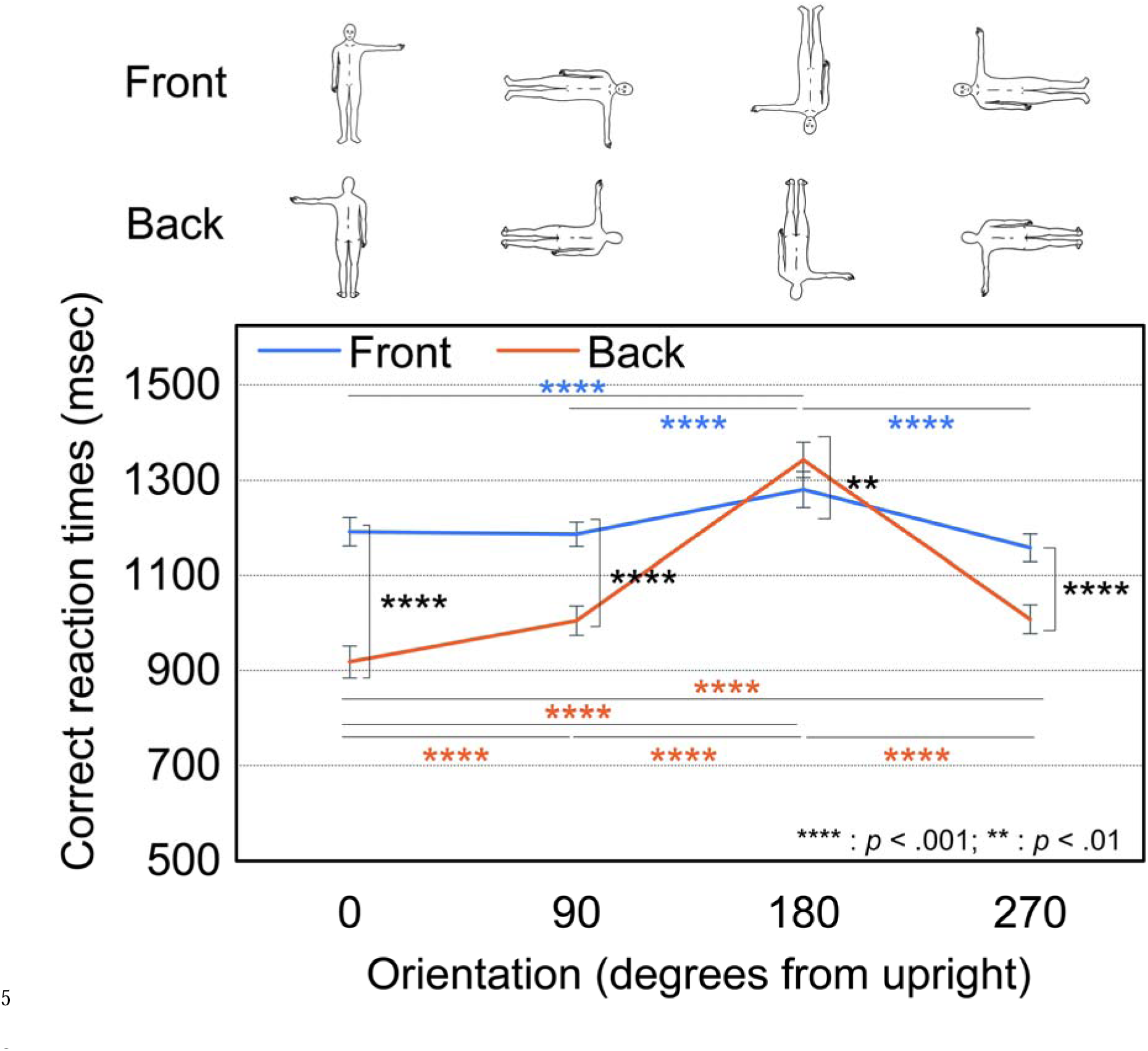
Correct reaction times (RT) for each stimulus. The error bars indicate the standard error of the mean.

#### 3.1.2. Correlation between correct reaction time and interoceptive sensitivity

To examine the relationship between task performance and individual differences in interoception, a correlation analysis was conducted between the IS and the ratio of correct RTs for front views to those for back views in each orientation. We found a significant negative correlation between the ratio of the correct RTs for the front views to those for the back views at 0° in the upright orientation and the IS (Skipped Spearman’s rho = -0.43887, *p* = 0.0336, adjusted for multiple comparisons using false-discovery rate (FDR) corrected, 95 % confidence interval = [-0.654571 - 0.137677]; Figure 3) after one outlier was excluded by using robust estimators (Pernet et al., 2013). This result indicates that individuals with a higher IS had a smaller ratio of correct RTs for front views than for back views in the upright orientation (0°).

**Figure 3.**
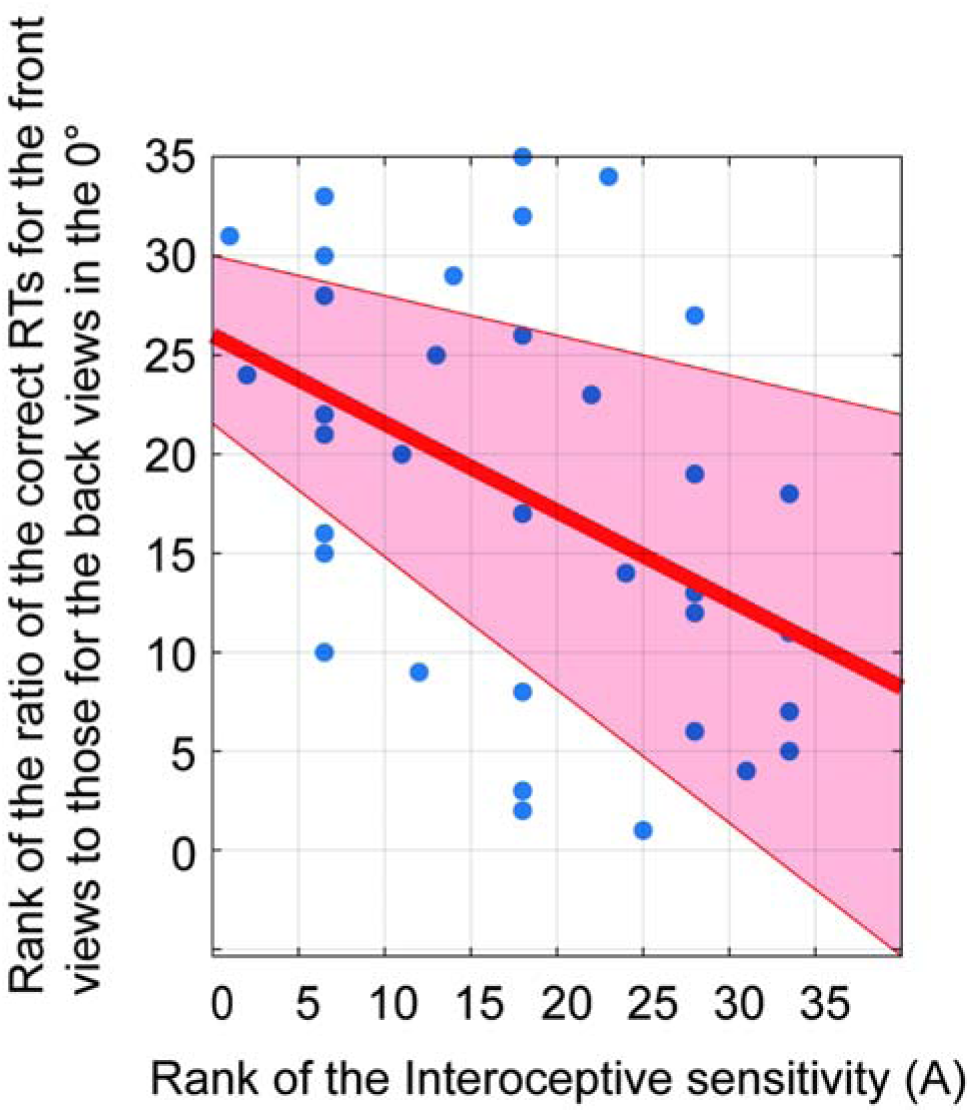
Correlation between interoceptive sensitivity, obtained through the heartbeat discrimination task, and the ratio of correct reaction times for the front views to those for the back views in 0° orientation. The 95 % confidence intervals are indicated by the red area. The solid line represents the result of the linear regression. RT, reaction time.

### 3.2. fMRI results

#### 3.2.1. Brain regions showing more activation for the front views than for the back views

Following the confirmation of the significant main effect of front/back through the two-way rmANOVA for the correct RTs, we examined the brain regions showing heightened activation for front views relative to back views. A comparison between brain activities for the front views and those for the back views revealed significant activations in the occipital and bilateral posterior occipitoparietal regions, including the superior parietal lobule (SPL), intraparietal sulcus (IPS), and frontal regions, including the left middle frontal gyrus and supplementary motor area (SMA). In addition, there were two clusters in the midline, including the regions from the medial superior frontal gyrus (mSFG) to the dACC extending to the pgACC, and the regions of the visual cortex and cerebellum (Figure 4 and Table 1).

**Figure 4.**
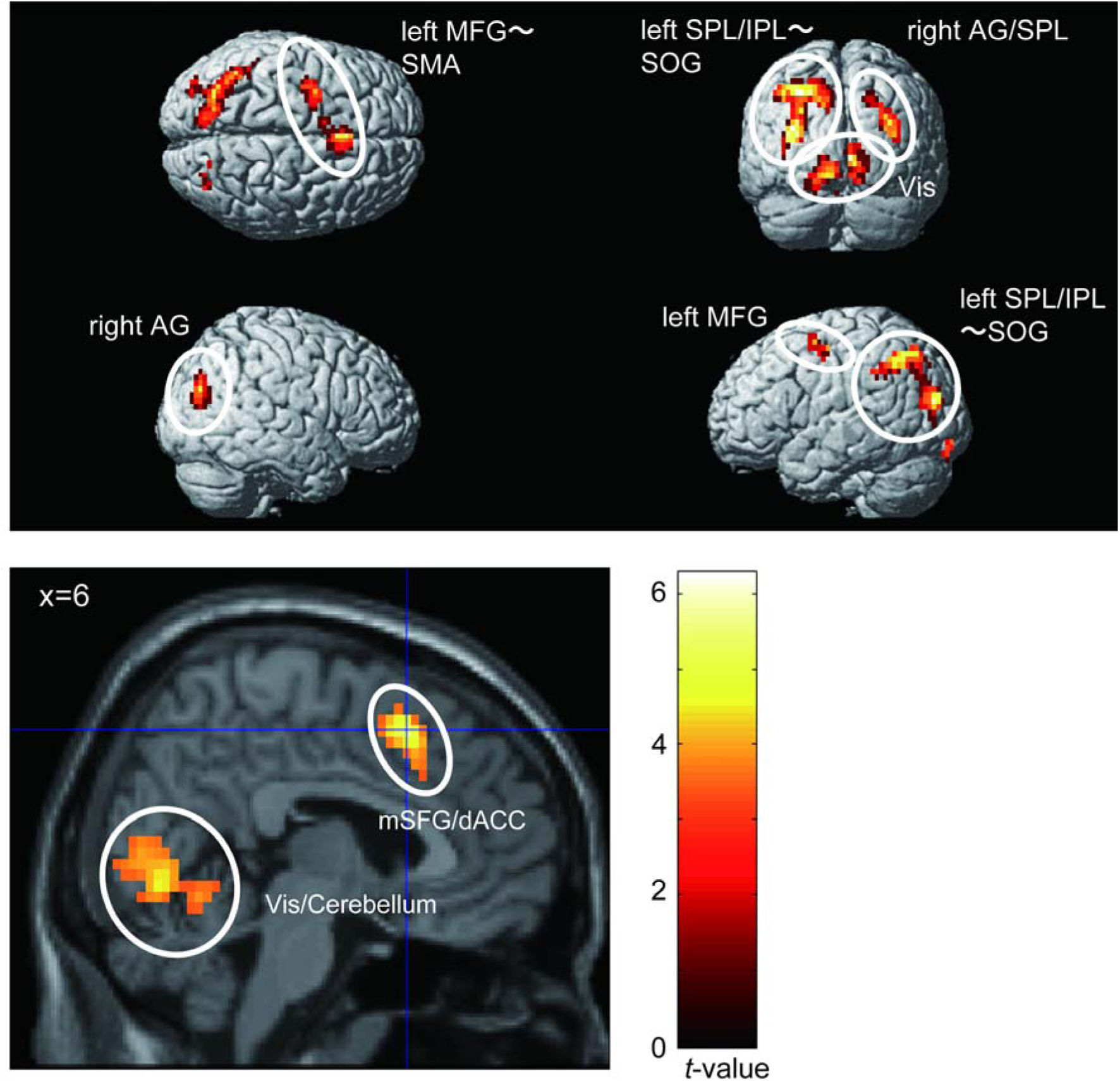
Brain regions showing more activation for the front view than for the back view during stimulus presentation. AG, angular gyrus; dACC, dorsal anterior cingulate cortex; IPL, inferior parietal lobule; MFG, middle frontal gyrus; mSFG, medial superior frontal gyrus; SMA, supplementary motor area; SOG, superior occipital gyrus; SPL, superior parietal lobule; Vis, visual area.

**Table 1.** Brain regions showing significantly higher activation for the front view compared to the back view during stimulus presentation.

| Anatomical region | Cluster<br>size | MNI coordinates<br>(mm) |  |  | *T*-value |
| --- | --- | --- | --- | --- | --- |
|  |  | x | y | z |  |
| Left superior parietal lobule | 1,504 | -33 | -61 | 42 | 6.26 |
| Left visual area |  | -12 | -79 | -2 | 6.16 |
| Left angular gyrus |  | -30 | -67 | 30 | 5.93 |
| Right medial superior frontal<br>gyrus | 256 | 6 | 17 | 50 | 5.39 |
| Left supplementary motor area |  | -27 | -4 | 54 | 8.41 |
| Left supplementary motor area |  | -9 | 5 | 54 | 4.24 |

3 Uncorrected $p < 0.001$ at the voxel level and family wise error-corrected $p < 0.05$ at the cluster level.
MNI, Montreal Neurological Institute.

The reverse comparison showed no significant regions, indicating that no region showed significantly more activation in the back views than in the front views. To examine the brain regions involved in the common processes for all stimuli, we conducted a conjunction analysis. This revealed three clusters: (1) a large cluster including the bilateral posterior regions extending from the occipital to the SPL and the parieto-motor regions extending from the bilateral temporo-parietal junction (TPJ) to the bilateral ventral/dorsal premotor and SMA; (2) the thalamus and brain stem; and (3) the right superior temporal gyrus (Supplementary Figure 1, Supplementary Table 1). To identify the brain regions involved in the transformation specific to front views, the brain regions showing more activation for front views than for back views were exclusively masked by the activation derived from the conjunction analysis. This procedure highlighted the activation of bilateral visual areas, the left SPL/inferior parietal lobule (IPL), and the mSFG/dACC (Supplementary Figure 2, Supplementary Table 2).

#### 3.2.2. Brain regions showing a similar trend to the correct reaction times

Based on the behavioral outcome that highlighted an interaction between front/back and orientation factors in the context of correct RTs, we also performed a two-way rmANOVA on brain activation during the stimulus presentations. This analysis incorporated the same factors of front/back and orientation. Activities in the mSFG/dACC and left dorsal premotor cortex (dPM) showed a significant interaction (Figure 5a and Table 2). The contrast estimates derived from the peak voxels within the mSFG/dACC and dPM clusters showed trends similar to those of the correct RTs for the front/back views and various orientations (Figure 5b).

**Figure 5.**
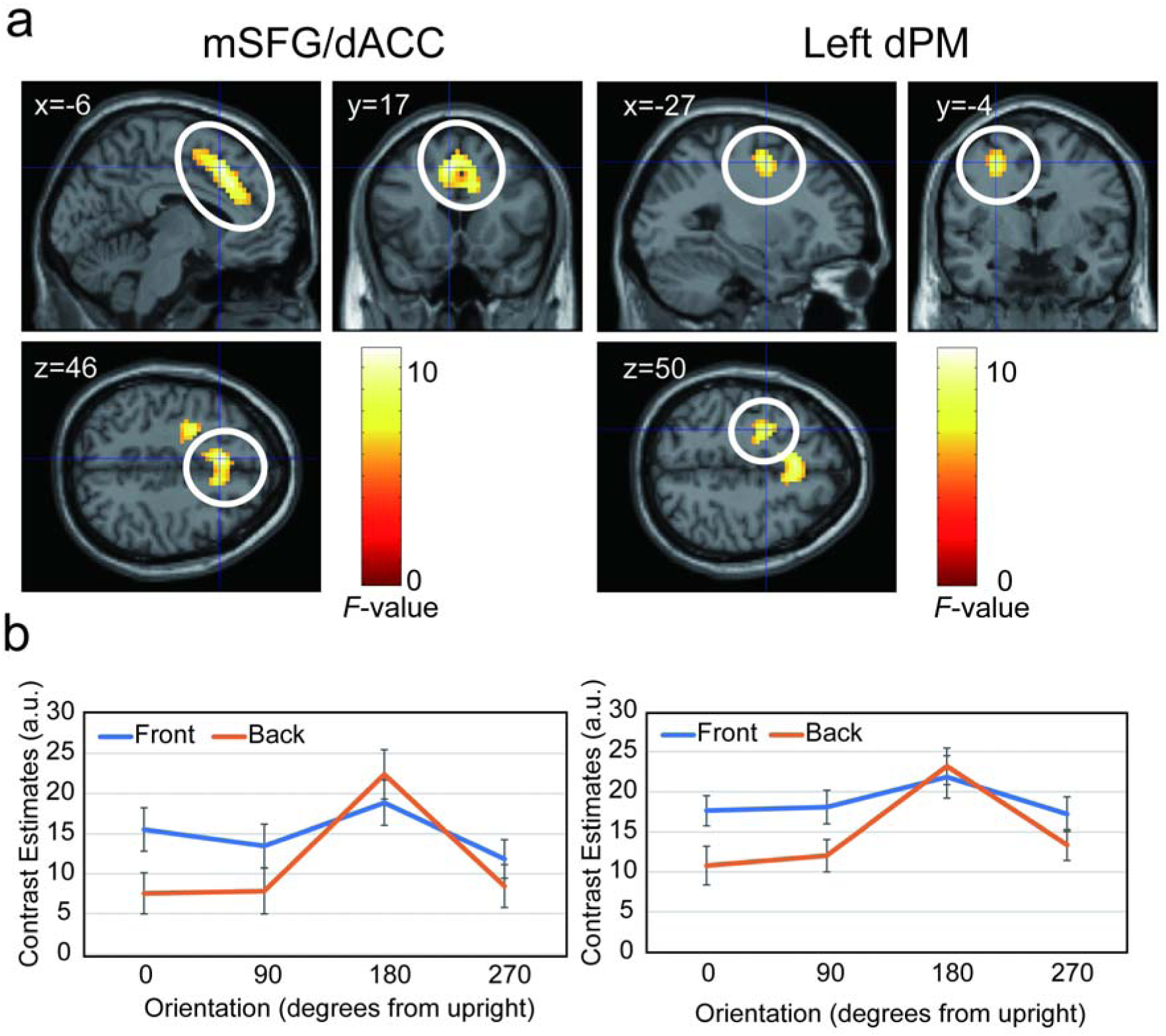
(A) Brain regions where activation showed an interaction of factors of front/back and orientation during stimulus presentation. (B) The contrast estimates of the peak voxel in the corresponding cluster shown in the upper panel for each view and orientation. dACC, dorsal anterior cingulate cortex; dPM, dorsal premotor cortex; mSFG, medial superior frontal gyrus.

**Table 2.**
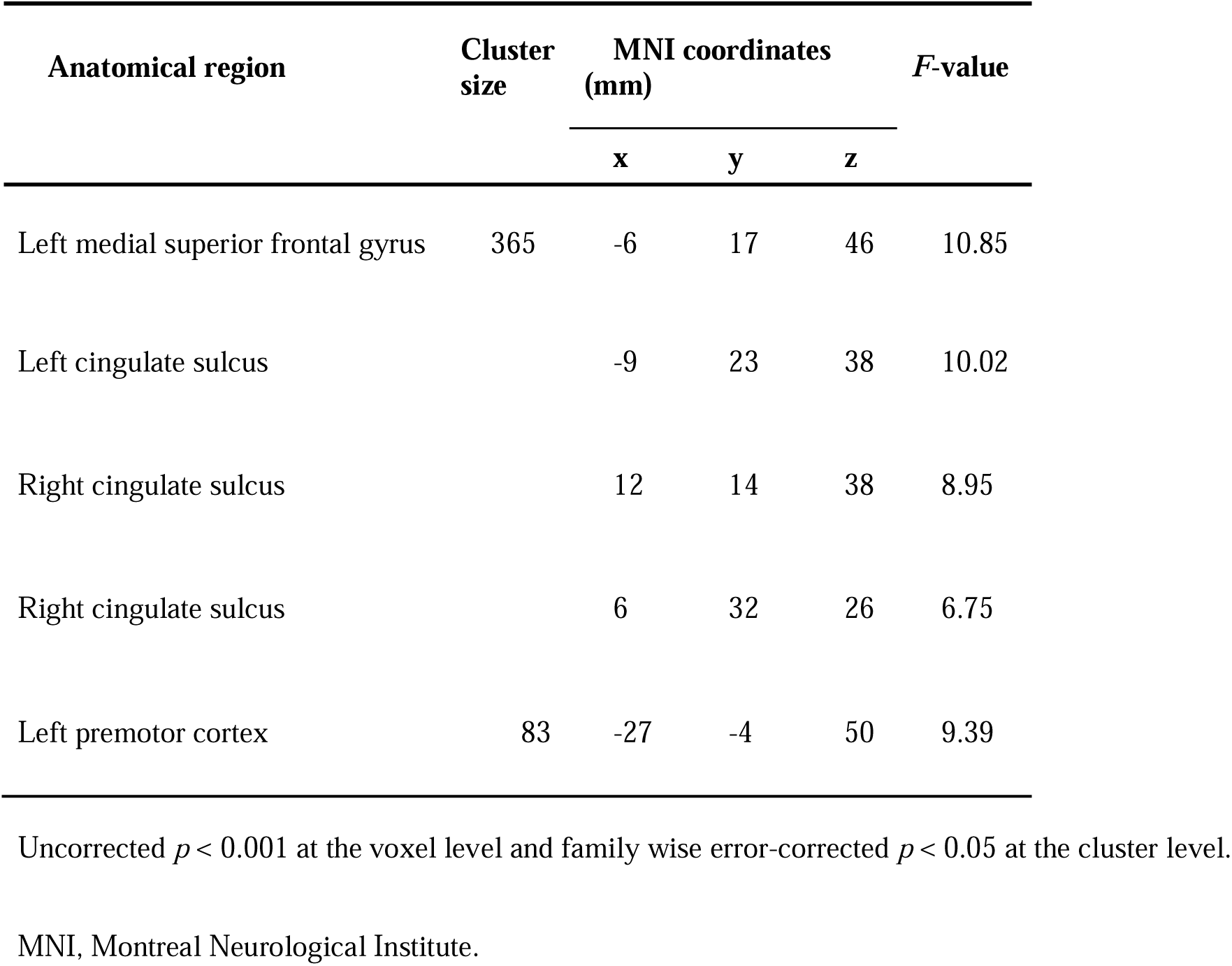
Brain regions showing a significant interaction between the factors of front/back and orientation during stimulus presentation.

#### 3.2.3. Brain activation related to the individual difference of interoceptive sensitivity

Following the finding that the activations within the mSFG/dACC and dPM clusters showed a similar trend to the correct RTs, we conducted a ROI analysis to examine whether these activations were associated with individual differences in IS. We defined nine anatomical ROIs in the bilateral mSFG (Brodmann areas (BA) 8 and BA 9), bilateral pgACC (BA 32), and left dorsolateral premotor area (BA 6) based on the regions included in the voxels of the mSFG/dACC and dPM clusters. The anatomical ROIs were defined using the Brainnetome atlas (Fan et al., 2016) as following: for the mSFG/dACC cluster, “A8m,” right/left medial area BA 8, “A9m,” right/left medial area BA 9, “cdACC,” right/left caudodorsal area BA 24, and “pgACC,” right/left pregenual area BA 32; for the dPM cluster, “A6dl,” left dorsolateral area BA 6, “A6vl,” left ventrolateral area BA 6, and “A6cdl,” left caudal dorsolateral area BA 6. We performed a correlation analysis using the contrast estimates of the front and back views averaged over the voxels in each ROI. This revealed that the mean contrast estimates in the bilateral medial area 8 (A8m), bilateral cdACC and pgACC, right medial area 9 (A9m), and left dorsolateral area 6 (A6dl) were positively correlated with IS (right A8m: Spearman’s rho = 0.476282, *p* = 0.020772; left A8m: Spearman’s rho = 0.409654, *p* = 0.025627; right cdACC: Spearman’s rho = 0.420195, *p* = 0.025627; left cdACC: Spearman’s rho = 0.470556, *p* = 0.020772; right pgACC: Spearman’s rho = 0.406141, *p* = 0.025627; left pgACC: Spearman’s rho = 0.364759, *p* = 0.040426; right A9m: Spearman’s rho = 0.363327, *p* = 0.040426; left A6dl: Spearman’s rho = 0.407442, *p* = 0.025627; all *p*s are adjusted for multiple comparison using FDR correction; Figure 6a).

**Figure 6.**
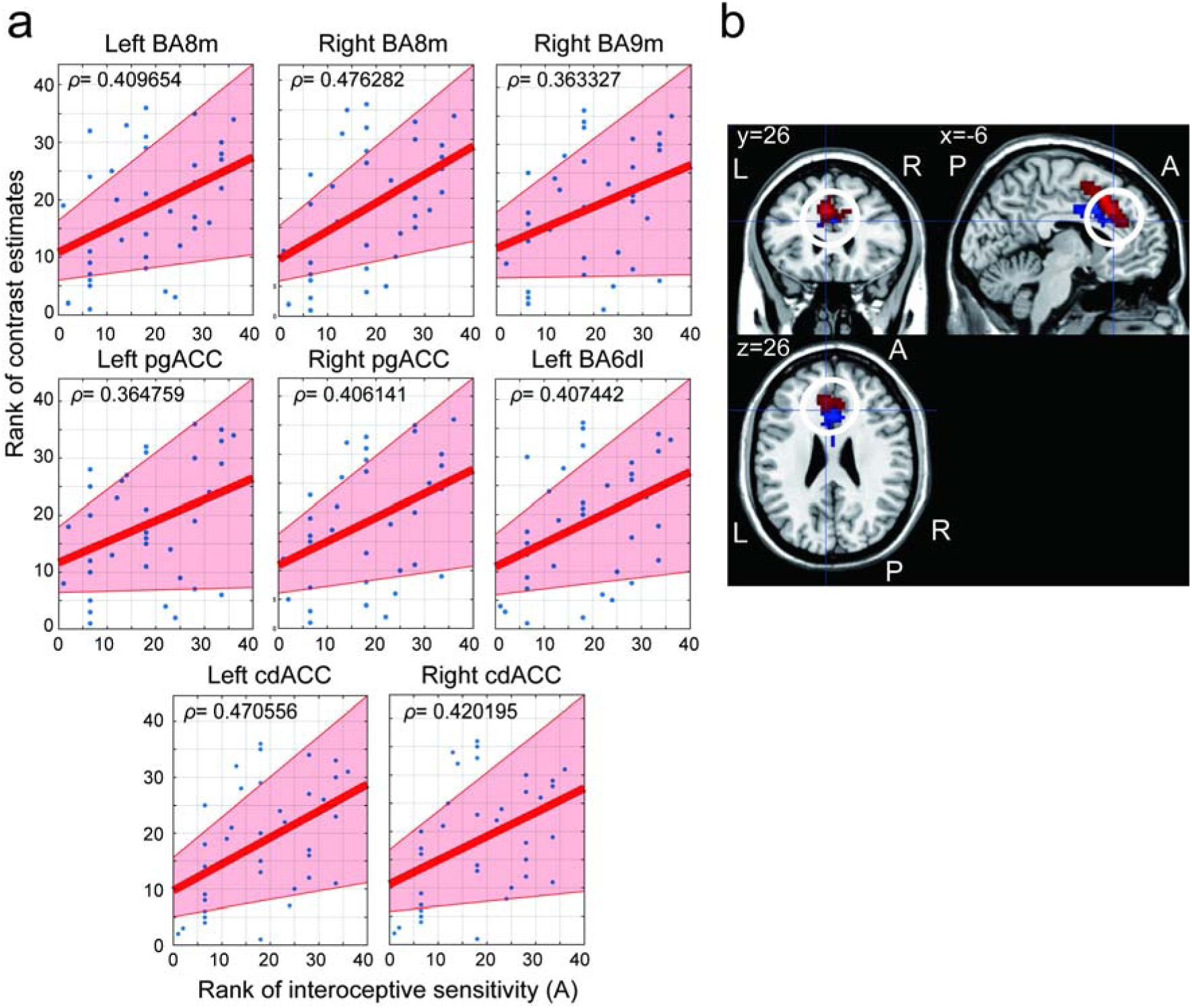
(A) Correlation between the interoceptive sensitivity of each participant derived from the heartbeat discrimination task and the contrast estimates averaged in each of clusters that showed a significant Spearman’s ρ. The solid line represents the result of a linear regression. The red area represents the 95 % confidence interval. (B) Brain regions where activation showed a significant correlation with the interoceptive sensitivity at a liberal statical threshold of uncorrected *p* < 0.005 at voxel level (FWE-corrected *p* = 0.0503 at cluster level; blue regions) and brain regions where activation showed an interaction of the factors of front/back and orientation in the cluster centered on the dACC (red regions). There was an overlap between them in the left pgACC (MNI cordinate x = -6, y = 26, z = 26). A, anterior; ACC, anterior cingulate cortex; BA6dl, dorsolateral Brodmann area 6; BA8m, medial Brodmann area 8; BA9m, medial Brodmann area 9; cdACC, caudodorsal anterior cingulate cortex; L, left; R, right. MNI, Montreal Neurological Institute; P, posterior; pgACC, pregenual anterior cingulate cortex.

We also performed a whole-brain regression analysis using IS as a covariate. This revealed a positive correlation between activations in the contrast (front > back) and interoceptive sensitivity in the dACC/pgACC cluster at a liberal statistical threshold of uncorrected *p* < 0.005 at the voxel level (FWE-corrected *p* = 0.0503 at cluster level). This cluster partly overlapped with the mSFG/dACC cluster, which showed an interaction between the front/back and orientation factors (Figure 6b, Table 3).

**Table 3.** Brain regions where the activations in the contrast (front > back) were positively associated with the interoceptive sensitivity.

| Anatomical region | Cluster size | MNI coordinates (mm) |  |  | T-value |
| --- | --- | --- | --- | --- | --- |
|  |  | x | y | z |  |
| Left dorsal ACC | 195 | -6 | 11 | 38 | 3.98 |
| Right ventral ACC |  | 0 | 20 | 26 | 3.47 |
| Right dorsal ACC |  | 9 | 14 | 34 | 3.43 |

#### 3.2.4. Functional connectivity

Based on the results that the cluster showed an interaction between the front/back and interaction factors, and that showed a positive association with the individual differences in interoception overlapped in the pgACC, we conducted a gPPI analysis to examine FC with the bilateral pgACC as the seed region. During the execution of ALJ tasks for front views, the left pgACC showed positive FC with bilateral clusters encompassing the pre/postcentral gyri, opercular part of the inferior frontal gyri, anterior supramarginal gyri, and posterior insulae (Figure 7a, Table 4a).

**Figure 7.**
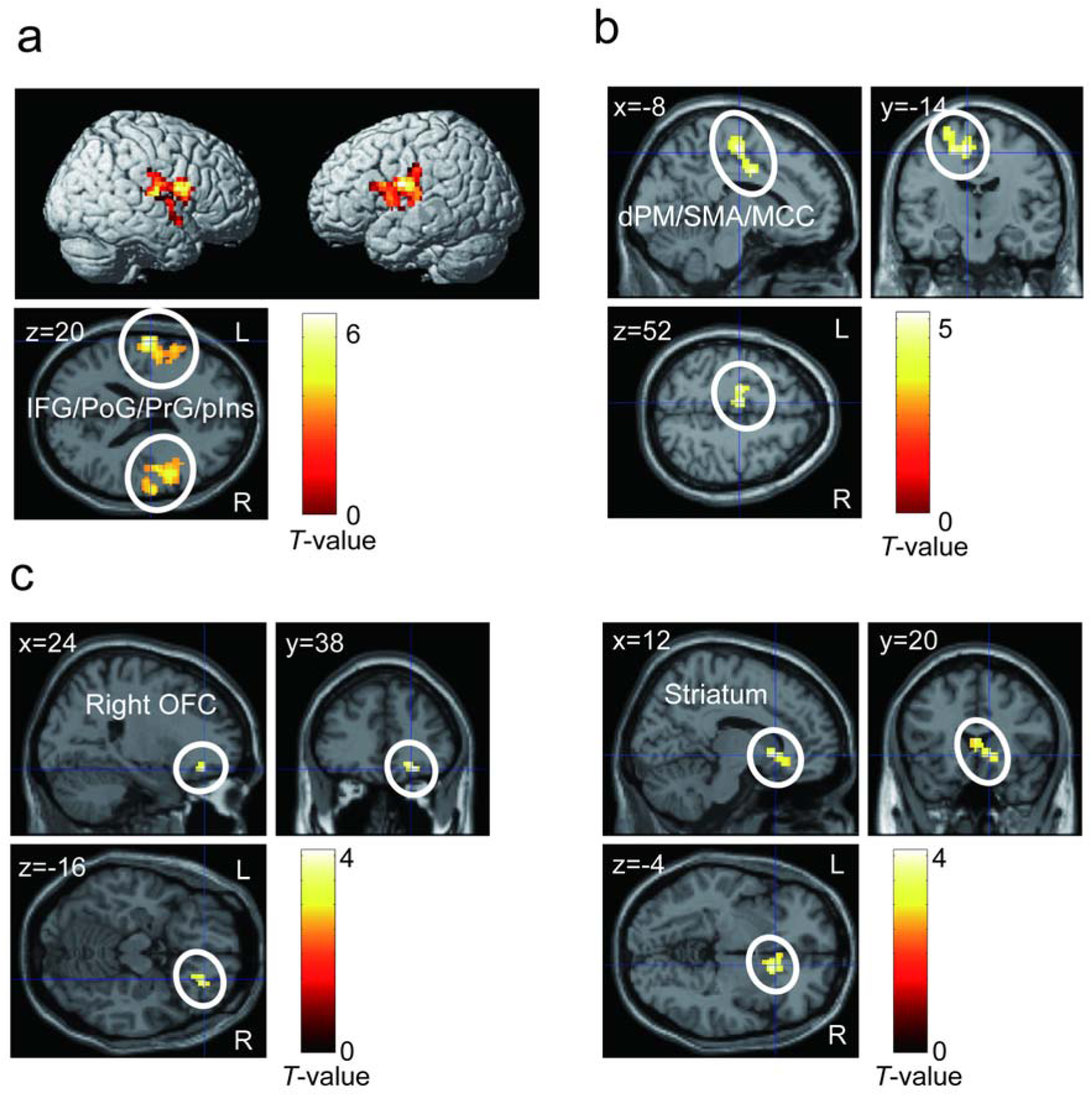
(A) Brain regions showing positive functional connectivity from the left pgACC during the presentation of the front views. (B) Brain regions showing positive functional connectivity from the left pgACC during the presentation of the front views. (C) Brain regions showing connectivity positively correlated with the individual difference in interoceptive sensitivity (IS) at a statistical threshold of uncorrected *p* < 0.005 at voxel level and false discovery rate-corrected *p* < 0.05 at cluster level. dPM, dorsal premotor area; IFG, inferior frontal gyrus; L, left; MCC, middle cingulate gyrus; OFC, orbitofrontal cortex; pIns, posteriror insula; PoG, postcentral gyrus, PrG; precentral gyrus; R, right.; SMA, supplementary motor area.

**Table 4.**
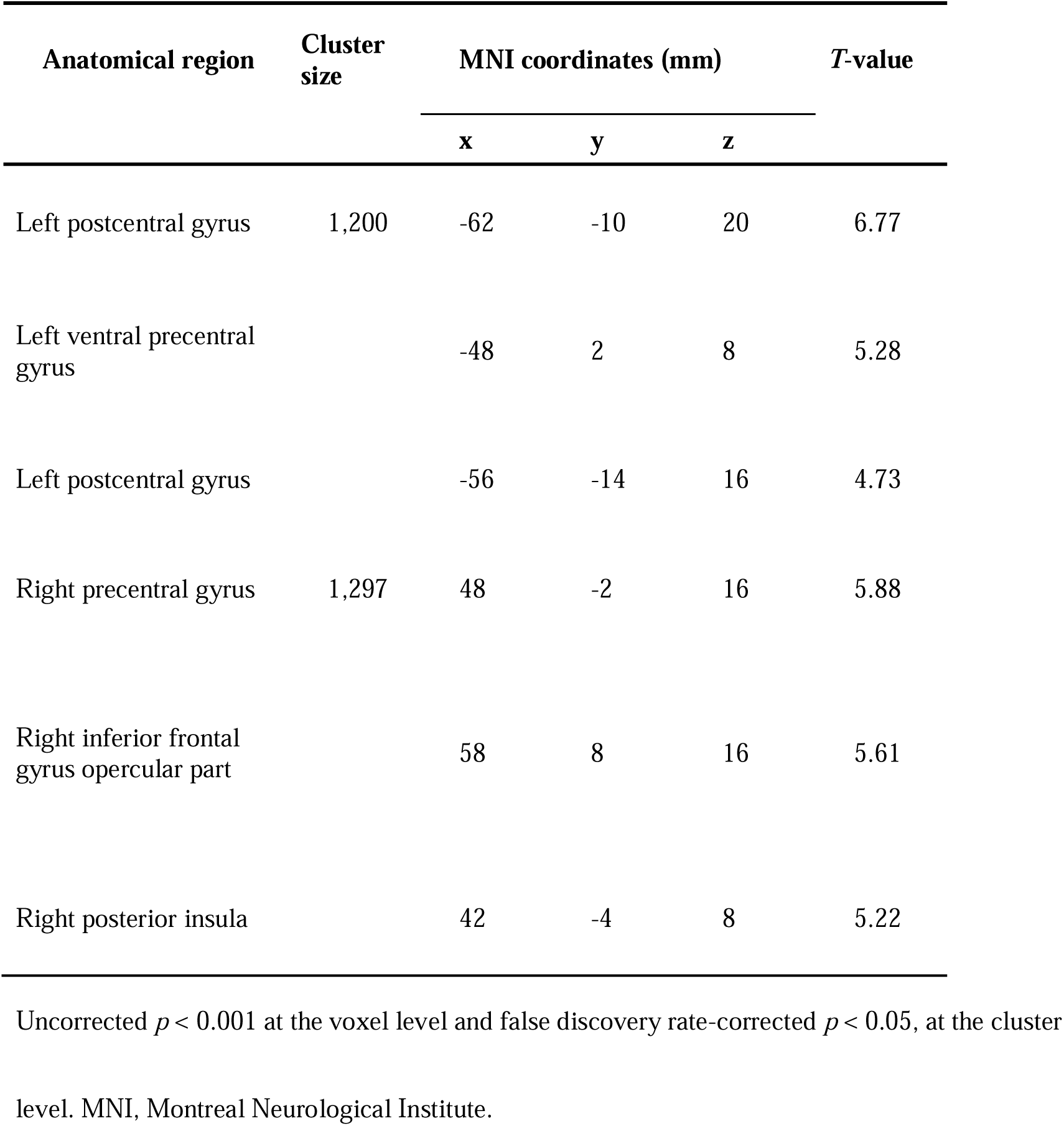
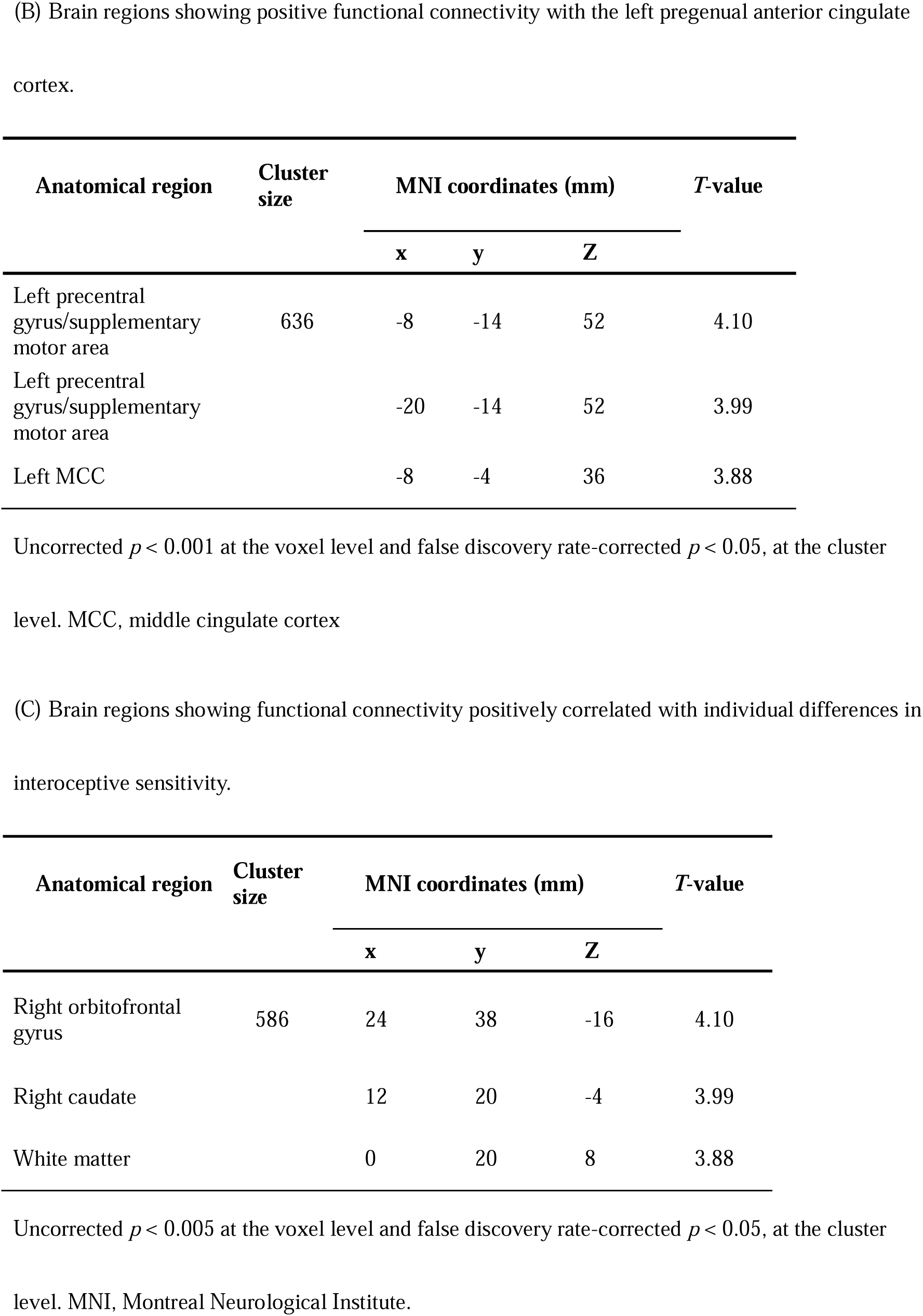
(A) Brain regions showing positive functional connectivity from the left pregenual anterior cingulate cortes.

| Anatomical region | Cluster size | MNI coordinates (mm) |  |  | *T*-value |
| --- | --- | --- | --- | --- | --- |
|  |  | *x* | *y* | *z* |  |
| Left postcentral gyrus | 1,200 | -62 | -10 | 20 | 6.77 |
| Left ventral precentral gyrus |  | -48 | 2 | 8 | 5.28 |
| Left postcentral gyrus |  | -56 | -14 | 16 | 4.73 |
| Right precentral gyrus | 1,297 | 48 | -2 | 16 | 5.88 |
| Right inferior frontal gyrus opercular part |  | 58 | 8 | 16 | 5.61 |
| Right posterior insula |  | 42 | -4 | 8 | 5.22 |
|  |  | **x** | **y** | **Z** |  |
| Left precentral gyrus/supplementary motor area | 636 | -8 | -14 | 52 | 4.10 |
| Left precentral gyrus/supplementary motor area |  | -20 | -14 | 52 | 3.99 |
| Left MCC |  | -8 | -4 | 36 | 3.88 |

| Anatomical region | Cluster size | MNI coordinates (mm) |  |  | *T*-value |
| --- | --- | --- | --- | --- | --- |
|  |  | **x** | **y** | **Z** |  |
| Right orbitofrontal gyrus | 586 | 24 | 38 | -16 | 4.10 |
| Right caudate |  | 12 | 20 | -4 | 3.99 |
| White matter |  | 0 | 20 | 8 | 3.88 |

- 7 Uncorrected $p < 0.005$ at the voxel level and false discovery rate-corrected $p < 0.05$ , at the cluster 8 level. MNI, Montreal Neurological Institute.

In contrast, the right pgACC showed a positive FC with a cluster that included the left precentral gyrus, SMA, and middle cingulate cortex (MCC) (Figure 7b, Table 4b). This left precentral area overlapped with the left dPM cluster, which showed the interaction of the front/back and orientation factors, as revealed by ANOVA (Supplementary Figure 3). To examine the FC related to individual differences in interoception, we explored the brain regions showing FC with the pgACC positively related to the IS when participants performed ALJ for front views in the upright orientation (0°). This revealed that the left pgACC showed positive FC with clusters including the striatum and right orbitofrontal cortex in a liberal statistical threshold of uncorrected *p* < 0.005 at the voxel level and FDR-corrected *p* < 0.05 at the cluster level (Figure 7c, Table 4c).

## 4. Discussion

In the present study, we tested three hypotheses that underpin the interplay between spatial image transformation of the self-body and interoception, using fMRI measurements throughout the execution of the ALJ task. H1, performance on the ALJ task correlates with individual differences in interoception, was validated by our behavioral data showing that the ratio of the correct RTs for the front views to those for the back views in the 0° orientation correlated positively with the amplitude of the Gaussian (IS) derived from the HDT. H2, the brain regions involved in exteroceptive imagery transformation and interoceptive processing are associated with performance on the ALJ task, which was also supported by our neuroimaging results showing that the activations in the dPM and the cluster including the mSFG/dACC exhibited trends similar to those of the correct RTs. H3, activities in the brain regions involved in the integration of intero-exteroceptive information correlated with individual differences in interoception during the ALJ task, were also validated by the results that activation in the cluster including the mSFG/dACC showed a significant positive correlation with the IS.

### 4.1. Behavioral data of ALJ

In the task used in the present study, which was identical to Parsons (1987) Experiment 1 albeit with a reduced stimulus set, the results were generally similar to those of Parsons (1987). For the back views, the RT was the longest in the 180° orientation, the RTs increased according to the clockwise/anticlockwise rotation angle from 0°. This pattern suggests the utilization of mental rotation mechanisms (Shepard and Metzler, 1971) in the execution of the ALJ for the back view stimuli. For the front views, the RT were longer than those for the back views, except in the 180° orientation. This discrepancy likely arose from the necessity to mentally transform and reconcile the disparities between the participant’s egocentric coordinate system and the visual representation of the body displayed in the stimuli, specifically concerning the left-right orientations. The RTs for front views were relatively independent of the orientation, suggesting that a combined transformation including the mental rotation to the upright orientation and the embodied simulation like “looking-back” was used to align the participant’s egocentric coordinate system with a presented stimulus. Parsons (1987) conducted an ALJ task in which line drawings of the body were presented in various orientations along various axes. The resultant rotation-dependent RTs for ALJ closely resembled the timing associated with an alternate task that required participants to mentally simulate the spatial transformation of their body to align with the stimulus orientation. This convergence between tasks suggested the involvement of physical simulation in the ALJ task. Therefore, our current findings not only replicated the outcomes of Parsons (1987) study within the MRI scanner but also provided further indications of embodied simulation’s engagement, specifically spatial image transformation of the self-body, in the execution of the ALJ task.

### 4.2. Brain activity specific to the front views

Within our fMRI dataset, we discerned augmented activation for front views compared to back views across several brain regions, encompassing the posterior parietal and dorsal premotor cortices, a cluster inclusive of the mSFG and dACC, and even the cerebellum. The engagement of the posterior parietal and dorsal premotor cortices have been frequently reported in tasks involving the mental imagery transformation (e.g., Hawes et al., 2019; Zacks, 2008). The cerebellum is known to have internal models that predict the sensory consequences of motor execution (Imamizu et al., 2000). Considering the previously proposed functions of the dACC, brain activation specific to front views suggests that the ALJ of front views may be performed by mental image transformation using motor imagery while monitoring bodily states. In contrast, no regions showed greater activation in the back views than in the front views. Moreover, in the analysis in which the contrast image for the comparison between the front and back views was exclusively masked with the conjunction for all stimuli, we found significant activation in the visual area, left IPS, and mSFG/dACC. These results suggest that there was no brain region that was specifically active in the processing of back views, but rather that there were brain regions engaging in general visual processing common to both front and back views, as well as brain regions specifically engaging in the processing of front views. These results are consonant with the theory that the ALJ task entails a combined transformation including mental rotation and “looking-back” component.

Moreover, we found that the activation of the clusters in the mSFG/dACC and left dPM showed an interaction between the factors of orientation and front/back, and the trends were similar to those of the correct RTs. Previous fMRI studies using hand laterality judgment (HLJ) have reported activation of the IPS and dorsal premotor cortex (e.g., Bonda et al., 1995; de Lange et al., 2006; Zapparoli et al., 2014). For instance, de Lange et al. (2006) reported left IPS activation when participants engaged in HLJ with hand drawings, especially when a considerable disparity existed between the participants’ hand posture and the posture of the depicted hand drawing. It has also been reported that the left IPS is active in cooperation with the left premotor cortex during visual-mental image transformation tasks (Sasaoka et al., 2014). Thus, the left IPS activity observed in the present study may reflect a visual image transformation linked to motor processing common to both the HLJ and ALJ. In contrast, the mSFG/dACC may be involved in processing related to the ALJ-specific looking-back transformation for front views. In the ALJ task, as in the HLJ task, the integration of multimodal information and efference copies from motor-related regions is important for performing the task. However, ALJ requires spatial image transformation of the entire body and generation of motor imagery while monitoring the interoception of the entire body. This may result in the activation of the cluster, including the dACC, which is involved in monitoring intero- and exteroceptive information, in addition to the left IPS.

### 4.3. Correlation between arm laterality judgment performance and interoceptive sensitivity

When the ratio of correct RTs for the front views to those for the back views was defined as the transformation cost to resolve the discrepancy with the egocentric coordinate system, the transformation cost at an ALJ of 0° for the front views showed a positive correlation with the IS. This correlation suggests a potential influence of individual differences in interoception on the embodied simulation used to align the egocentric coordinate system with the presented stimulus.

Regarding the relationship between interoception and the sense of self, Tsakiris et al. (2011) showed that participants with lower interoceptive accuracy, as determined by a heartbeat counting task, reported greater proprioceptive drift in the RHI. However, the relationship between the amount of RHI and interoception has been less clear in other studies (e.g., Crucianelli, et al., 2018). In contrast, Suzuki et al. (2013) conducted an experiment in which a participant’s hand was presented in virtual reality at a different position from their actual hand and the RHI was elicited by cardiac feedback projected on the virtual hand or tactile stimulation. They reported a positive association between the RHI elicited by tactile stimulation on participants’ hands and the IS derived from the HDT. Thus, the relationship between individual differences in interoception and the occurrence of detachment from the self-body has yielded inconsistent outcomes across different studies.

One reason for this inconsistency is the difference in the tasks used in these studies to assess individual differences in interoception (Suzuki et al., 2013). The HDT can be considered a task for assessing individual differences in intero-exteroceptive integration. When multiple exteroceptive stimuli (e.g., visual and tactile stimuli) are presented consistently, individuals who can efficiently integrate intero-exteroceptive information may be able to associate them with their interoception in a flexible manner. This may result in a positive correlation between the performance of the ALJ and the IS derived from the HDT.

The absence of correlations between IS and ALJ performance for the orientations, except at 0° might be caused by the following reasons: if the two kinds of transformations, looking-back and mental rotation, are involved in the ALJ, the ALJ of the front views in the 0° orientation would require a looking-back transformation only, whereas the ALJ for the front views in other orientations would require a combined transformation of looking-back and mental rotation. Individual differences in IS were evident for the ALJ in the 0° orientation because the looking-back transformation employed an embodied simulation. In contrast, other factors of individual differences, such as visual working memory (WM) capacity (Zhang et al., 2022), might affect mental rotation or a combination of the two types of transformations. Accordingly, these results support the involvement of interoceptive processing in embodied simulation.

### 4.4. Brain activations and functional connectivity associated with the interoceptive sensitivity

The ROI analysis, based on the brain regions where activations showed an interaction of the front/back and orientation factors, unveiled predominantly positive correlations with IS. In the whole-brain analysis, a cluster within the dACC/pgACC exhibited a positive correlation with IS, albeit at a liberal statistical threshold. This cluster overlapped with the pgACC region where activations showed an interaction, these results indicate that the pgACC is a region involved in the ALJ associated with individual differences in interoception.

The index for individual differences in interoception used in the present study was obtained by curve fitting the participants’ responses to multiple delay time conditions in the HDT. Based on signal detection theory, the amplitude of the Gaussian curve (IS) can be interpreted as an index that expands as the signal-to-noise ratio of heartbeat perception is higher. Based on the free-energy principle (Friston, 2010), it can be speculated that individuals with high IS correspond to those with high prediction error accuracy. Maekawa et al. (2021) reported that participants’ heart rates increased in response to music stimuli that they rated as high valence, and that activation in the middle insula was more evident in the high IS group than in the low IS group. These results suggest that individuals with higher IS are more sensitive to the physiological changes that occur in response to musical stimuli and that these changes are reflected in middle insular activity. Similarly, many previous studies have reported a relationship between individual differences in interoception and insular activity (e.g., Critchley et al., 2004, Tan et al., 2018, Haruki et al., 2021). In the present study, however, the IS demonstrated no discernible link with insula activity; instead, the ACC belonged to the salience network as the insula. It has been proposed that the ACC monitors the internal and external environments (Heilbronner and Hayden, 2016). Given this hypothesis, a discrepancy between intero-exteroceptive states was detected in the ACC. In individuals with high IS, the precision of discrepancy detection is higher than that in individuals with low IS, resulting in ACC, especially pgACC, activity. Additionally, the fact that the dPM also showed a positive correlation with the IS might be caused by the positive FC to the dPM from the right pgACC as the seed region.

The fact that we observed no relationship between IS and insular activity could lead to the following interpretations: Most previous research has shown a relationship between the insular cortex and individual differences in interoception mainly in emotional processing. However, the ALJ task does not require emotional processing; rather, it engages cognitive-motor processing. Within the salience network, the anterior insula can be considered a ‘limbic sensory area’ and the ACC a ‘limbic motor area’ (Seth, 2013), and the ACC is considered to be involved in the initiation of behavior based on detected salience (Craig, 2009). For these reasons, we did not observe any involvement of the insula in this study.

The gPPI analysis using the pgACC as a seed region showed that during the presentation of the front views, the left pgACC showed FC to the bilateral pre/postcentral gyri, anterior inferior supramarginal gyri, and posterior insula, whereas the right pgACC showed FC to the MCC and SMA, some of which overlapped with the dPM region where interactions were observed. The regions that showed FC from the left pgACC included the parieto-insular vestibular region (e.g., Bottini et al., 1994) and the posterior insula, which receives vestibular and interoceptive information (e.g., Craig, 2009), as well as the anterior TPJ, which is reportedly involved in VPT (e.g., Ganesh et al., 2015). In particular, the TPJ has been suggested to be involved in the computation of the prediction error in terms of free-energy theory (Doricchi et al., 2022). Through multisensory integration, including visual, somatosensory, vestibular, and interoceptive information, implemented by the neural network among these regions and the pgACC, it can be speculated that ALJ is performed by the spatial imagery transformation of one’s body based on the computation of errors using the difference between the postures of the actual body and the presented stimulus.

Moreover, the left pgACC showed interoception-related FC with a cluster including the ventral striatum, such as the nucleus accumbens and the OFC. The nucleus accumbens is known as “limbic-motor interface” (Mogenson et al., 1980) to transfer motivation into motor responses. The striatum receives interoceptive ascending projections from the viscera, which underpin behavioral changes (Savitz and Harrison 2018). Considering that the ACC is a limbic motor region, the present results suggest that the pgACC plays an important role in the integration of intero-exteroceptive information on the body for the embodied simulation of the spatial transformation of mental body imagery in cooperation with the premotor cortex.

The mSFG/dACC cluster observed in this study also overlapped with the rostral cingulate zone (RCZ; Shackman et al., 2011). The RCZ is commonly referred to cingulate motor area (Vogt et al., 2003), and its subregions are organized somatotopically. For example, the facial region controls the muscles of the upper face. This suggests that the RCZ is involved in expressing affect and executing goal-directed behaviors. The RCZ has been identified in humans with BAs 32’ and a24c’ in the vicinity of the cingulate sulcus. In the somatotopic map shown by Schackman et al., our local peak coordinates of the cluster in the cingulate sulcus (6, 32, 26), where the activations showed that the interaction corresponded to the arm region. Moreover, it has been reported the RCZ neurons receive interoceptive information through the spinothalamic tract via the mediodorsal thalamus (Shackman et al., 2011). Although the RCZ is classified as the MCC, the mSFG/dACC cluster observed in the present study extended anteriorly in the medial areas along the RCZ, extended to the dACC, and reached the pgACC. In particular, the pgACC correlated with the IS, suggesting that integration with interoception occurs more anteriorly and that motor imagery recruits the RCZ and premotor area. This is consistent with the recently proposed information pathway for action-outcome learning from the ACC to the midcingulate motor area and then to the premotor areas, including the SMA (Rolls, 2019).

Furthermore, the gPPI analysis revealed different brain regions showing FC with the left and right pgACC, suggesting laterality in ACC function. For instance, in an fMRI study using a go-no-go task (Lütcke and Frahm, 2008), based on the finding that the right ACC was active when participants correctly inhibited their response, they suggested that the right ACC is involved in cognitive control and that the bilateral ACC is involved in error processing. These results are consistent with our finding that the right pgACC showed FC with the SMA and dPM.

However, another interpretation of the function of the ACC in the ALJ is possible, since the ACC has been reported to be involved in visual WM (e.g., Petit et al., 1998; Courtney et al., 1996; Mesulam et al., 2001) and attentional control (e.g., Luks, et al., 2002; Langner and Eickhoff, 2013) and the peak voxel of the dPM cluster was observed in the frontal eye field (e.g., Linden, et al., 2003), which may reflect the influence of visual WM and attention. A meta-analysis of the delay activity during visual WM (Li et al., 2023) also showed activation of the MCC (BAs 24 and 32), which seems to be spatially close to the posterior part of the mSFG/dACC cluster found in the present study, suggesting that a part of this cluster reflects attention-related activities. However, the absence of activity in the dorsolateral prefrontal cortex, which is generally considered to be involved in WM and attention control, and the positive correlation between mSFG/dACC activity and IS in the processing of the upright view that required no mental rotation, which is considered to have a relatively small WM load. This suggests that the results of this study cannot be explained by the effects of attention or visual WM alone.

### 4.5. Limitations and future directions

Although proprioception accuracy is expected to influence the spatial image transformation needed for the ALJ task, it was not examined in this study. Proprioception is sometimes classified as interoception (Dieter, 1996). Moreover, proprioception, but not interoception, is associated with the occurrence of RHI (Horváth et al., 2020), future studies should distinguish between proprioception and interoception and examine the effect of individual differences on ALJ performance. In addition, differences in vestibular sensation among individuals may be involved in the generation of accurate rotational images of the body, and may need to be examined separately from individual differences in interoception. Therefore, future studies should clarify the relationship between the following three factors that potentially affect the spatial image transformation of the self-body: individual differences in interoception, vestibular sensation, and proprioception. This provides further insights into the mechanisms of human self-body perception and body schemas.

In the present study, the ALJ task was performed while participants were lying down inside an MRI scanner, whereas for most behavioral experiments, participants were seated in a chair. In particular, in the 180° front view scenario, the most direct way to align the participants’ body with the stimulus is by leaning backwards; however, it may be easier to create such motor imagery in the lying posture than in the sitting posture. This distinction could potentially lead to a reversal in the correct RT between the back and front views for the 180° orientation. The influence of posture on the ALJ performance should be investigated in future studies.

## 5. Conclusions

In the present study, we demonstrated an association between individual differences in interoception and spatial image transformation of the self-body, using the ALJ task. Our neuroimaging findings indicate the central role of the ACC in integrating the intero-exteroceptive information necessary for executing the ALJ task. These results provide the first instance of neurobehavioral evidence of the involvement of interoceptive processing in the complex cognitive process of transforming the spatial representation of the self-body. Furthermore, we demonstrated the engagement of ACC in the cognitive-motor functions related to interoceptive processing. This stands in contrast to the previously reported role of the anterior insula. Thus, these findings offer a fresh perspective on the neural mechanisms governing the processing of interoceptive and self-related information processing.

## CRediT authorship contribution statement

**Takafumi Sasaoka:** Conceptualization, Methodology, Formal analysis, Funding acquisition, Writing – original draft, Visualization. **Kenji Hirose:** Software, Formal analysis, Investigation, Writing – Review & Editing. **Toru Maekawa:** Methodology, Software, Writing – Review & Editing. **Toshio Inui**: Conceptualization, Supervision, Project administration, Funding acquisition, Writing – Review & Editing. **Shigeto Yamawaki:** Supervision, Funding acquisition, Writing – Review & Editing.

## Acknowledgments

This work was supported by Grants-in-Aid for Scientific Research from the Japan Society for the Promotion of Science (Grant No. 20K20423) and the Japan Science and Technology Agency (JST) COI (Grant Nos. JPMJCE1311 and JPMJCA2208), and the JST Moonshot Program Goal 9 (Grant No. JPMJMS2296). We are grateful to Dr. Nobuhiko Asakura, Dr. Kenji Ogawa, Dr. Alan S. R. Fermin, Dr. Kentaro Ono, and Dr. Maro G. Machizawa for helpful discussions concerning this study, and Ms.

Ayumi Uchida and Ms. Tamami Tomita for assistance with the implementation of the experiment. We would like to thank Editage (www.editage.com) for the English language editing.

## Data availability statements

The data supporting the conclusions of this manuscript will be made available by the authors without undue reservation.

