## Supplemental Figures and Tables for "The anterior cingulate cortex is involved in intero-exteroceptive integration for spatial image transformation of the self-body"

**Supplementary Materials**

### 1 Supplementary Figure

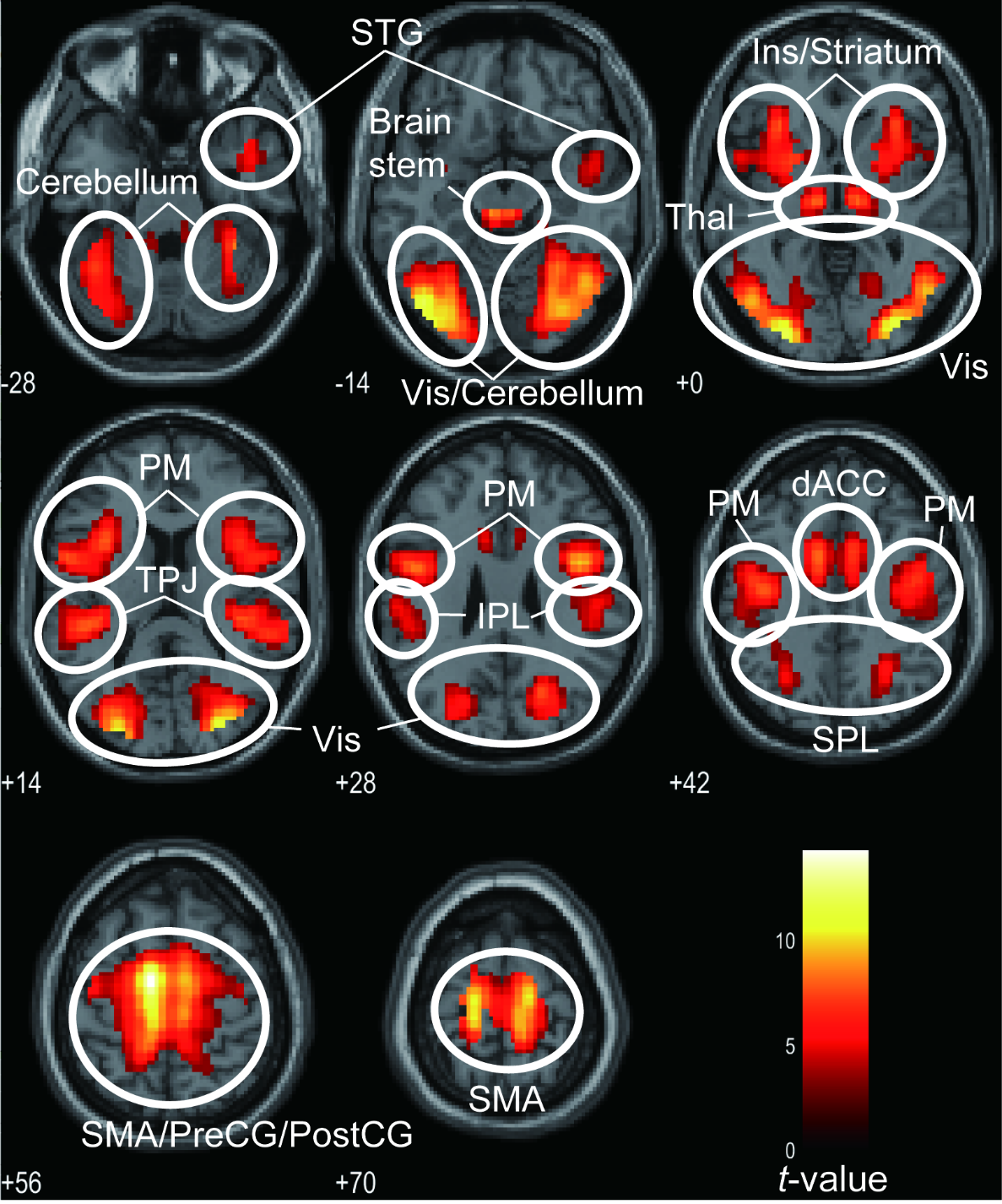
 **Supplementary Figure 1.** The result of the conjunction analysis. dACC, dorsal anterior cingulate cortex; Ins, insula; IPL, inferior parietal lobule; PM, premotor cortex; PostCG, postcentral gyrus; PreCG, precentral gyrus; SMA, supplementary motor area; SPL, superior parietal lobule; STG, superior temporal gyrus; Thal, thalamus; TPJ, temporo-parietal junction; Vis, visual area.

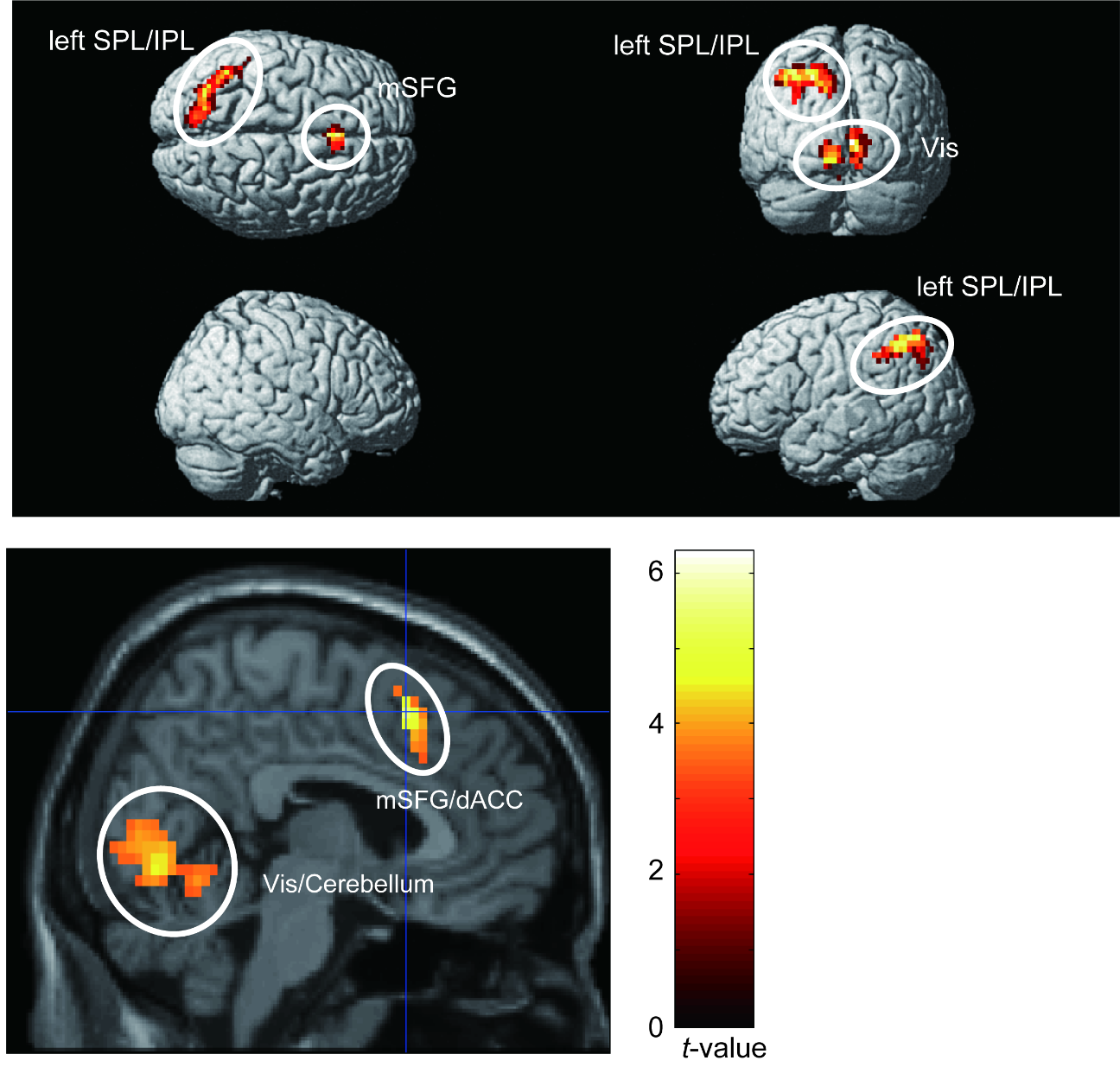

**Supplementary Figure 2.** The brain regions, that showed more activation for the front views than for the back views, exclusively masked with the activation revealed from the conjunction analysis. dACC, dorsal anterior cingulate cortex; IPL, inferior parietal lobule; mSFG, medial superior frontal gyrus; SPL, superior parietal lobule; Vis, visual area.

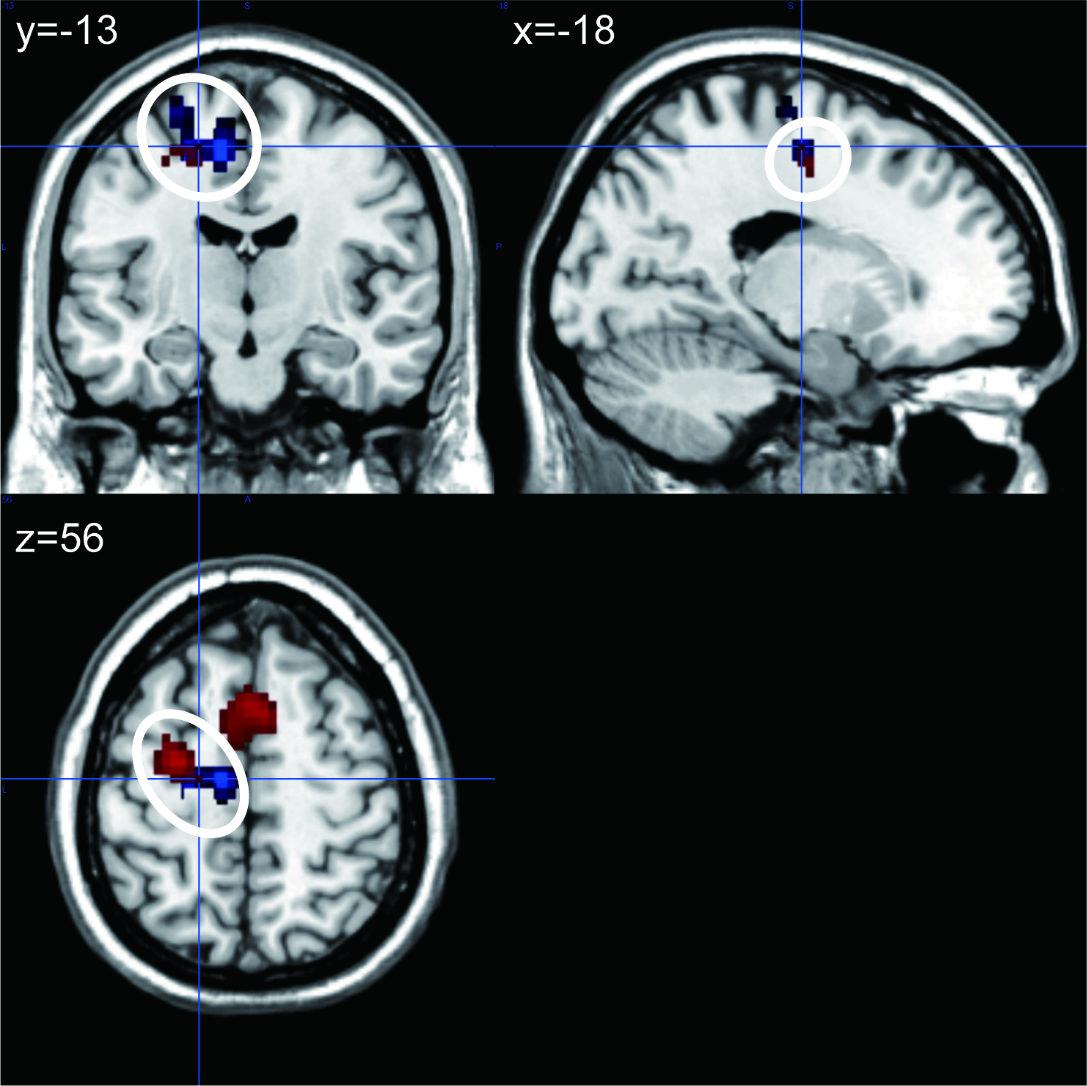

**Supplementary Figure 3.** The overlap between the left precentral area that showed a positive FC with the right pgACC (blue regions) and the left dorsal premotor cluster that showed an interaction of the factors of front/back and orientation revealed by ANOVA (red regions).

### 2 Supplementary Table

**Supplementary Table 1.** The results of the conjunction analysis.

| **Anatomical region** | **Cluster size** | **MNI coordinates (mm)** | | | ***T*-value** |
| --- | --- | --- | --- | --- | --- |
|  |  | **x** | **y** | **Z** |  |
| White matter | 8,281 | -9 | -10 | 58 | 14.31 |
| Left visual area |  | -45 | -76 | -10 | 12.74 |
| White matter |  | -15 | -28 | 66 | 12.59 |
| Brainstem | 304 | -6 | -22 | -10 | 9.02 |
| Brainstem |  | 6 | -22 | -10 | 8.97 |
| Left thalamus |  | -9 | -19 | -2 | 8.82 |
| Right superior temporal gyrus | 124 | 45 | 2 | -22 | 6.09 |
| Right temporal pole |  | 42 | 11 | -26 | 5.42 |

Uncorrected *p* < 0.001 at the voxel level and family wise error-corrected *p* < 0.05, at the cluster level. MNI, Montreal Neurological Institute.

**Supplementary Table 2.** The brain regions that showed more activation in front views than in back views were exclusively masked by the activation revealed by conjunction analysis.

| **Anatomical region** | **Cluster size** | **MNI coordinates (mm)** | | | ***T*-value** |
| --- | --- | --- | --- | --- | --- |
|  |  | **x** | **y** | **Z** |  |
| Left superior parietal lobule | 856 | -33 | -61 | 42 | 3.98 |
| Left visual area |  | -12 | -79 | -2 | 3.47 |
| White matter |  | -30 | -64 | 26 | 3.43 |
| Right medial superior frontal gyrus | 83 | 6 | 17 | 50 | 5.42 |

Uncorrected *p* < 0.001 at the voxel level and family wise error-corrected *p* < 0.05, at the cluster level. MNI, Montreal Neurological Institute.
